# Transcriptomic characterization of recombinant *Clostridium beijerinckii* NCIMB 8052 expressing methylglyoxal synthase and glyoxal reductase from *Clostridium pasteurianum* ATCC 6013

**DOI:** 10.1101/2024.02.27.582317

**Authors:** Santosh Kumar, Eric Agyeman-Duah, Victor C. Ujor

## Abstract

Bioconversion of abundant lactose-replete whey permeate to value added chemicals holds promise for valorization of this increasing food processing waste. Efficient conversion of whey-permeate-borne lactose requires adroit microbial engineering to funnel carbon to the desired chemical. Having engineered a strain of *Clostridium beijerinckii* NCIMB 8052 (*C. beijerinckii*_mgsA+mgR) that produces 87% more butanol on lactose than the control strain, in this study, we deployed RNA sequencing to profile the global transcriptome of *C. beijerinckii*_mgsA+mgR. The results revealed broadly contrasting gene expression patterns in *C. beijerinckii*_mgsA+mgR relative to the control strain. These were characterized by widespread downregulation of Fe-S proteins in *C. beijerinckii*_mgsA+mgR, coupled with increased expression of lactose uptake and catabolic genes, iron and phosphate uptake genes, two component signal transduction and motility genes, and genes involved in the biosynthesis of vitamin B_5_ and B_12_, aromatic amino acids, particularly tryptophan; arginine, and pyrimidines. Conversely, L-aspartate-dependent *de novo* biosynthesis of NAD as well as biosynthesis/metabolism of glycine, threonine, lysine, isoleucine and asparagine were downregulated. Furthermore, genes involved in cysteine and methionine biosynthesis and metabolism, including cysteine desulfurase—a central player in Fe-S cluster biosynthesis—were equally downregulated. Genes involved in biosynthesis of capsular polysaccharides and stress response were also downregulated in *C. beijerinckii*_mgsA+mgR. The results suggest that remodeling of cellular and metabolic networks in *C. beijerinckii*_mgsA+mgR to counter likely effect of methylglyoxal production following heterologous expression of methyl glyoxal synthase led to enhanced growth and butanol production in *C. beijerinckii*_mgsA+mgR.

**IMPORTANCE:** Biological production of commodity chemicals from abundant waste streams such as whey permeate represents a rational approach for decarbonizing chemical production. Whey permeate remains a vastly underutilized feedstock for bioproduction purposes. Thus, enhanced understanding of the cellular and metabolic repertoires of lactose-mediated production of chemicals such as butanol, promises to arm researchers with new engineering targets that can be fine-tuned in recombinant and native microbial strains to engender stronger coupling of whey permeate-borne lactose to value-added chemicals. Our results highlight new genetic targets for future engineering of *C. beijerinckii*_mgsA+mgR and indeed, *C. beijerinckii* for improved butanol production on lactose, and ultimately in whey permeate.

## INTRODUCTION

With increasing global demand for cheese, the generation of whey—a byproduct of cheese and casein production—is expected to continue to increase significantly. According to Kosikowski (1), ∼9 kg of whey is generated per kg of cheese produced. Global cheese production was predicted to reach 25.3 million metric tonnes in 2023 (2), which translates to the accumulation of 227.7 metric tonnes of whey. The dairy industry deploys ultrafiltration to separate the protein content of whey, which is sold as a supplement (3–5). This leaves behind a lactose-rich effluent referred to as whey permeate (WP), which places a considerable economic burden on the industry, whilst posing a hazard to the environment (4–6). With biochemical oxygen demand and chemical oxygen demand ranging from 40,000–48,000 mg/L and 80,000–95,000 mg/L, respectively (4,7), land spreading of WP constitutes a major hazard to surface water bodies (4–6). Although WP is used to compound livestock feed, this practice is dwindling, thus encumbering the dairy industry with WP treatment costs. Spray-drying to produce whey powder attracts additional costs, which makes this approach economically less attractive (4,8). Consequently, developing cost effective strategies for the valorization of WP promises considerable economic and environmental payoffs.

Towards valorization of WP, we sought to engineer *Clostridium beijerinckii* NCIMB 8052 (hereafter *C. beijerinckii*) to co-produce 1,2-propanediol (1,2-PDO) and butanol on lactose (WP sugar). Both 1,2-PDO and butanol are value-added chemicals with multifarious applications across the economy, and are currently produced solely from fossil fuel feedstocks. Butanol has enormous potential as renewable fuel for aviation and automobile applications, as a feedstock in the production of plasticizers, synthetic rubber, paints, and as a broad-application industrial solvent (9–11). Similarly, 1,2-PDO finds applications in the manufacture of polyester resins, building materials, non-ionic detergents, and cosmetics, and as a pharmaceutical solvent (12–15). Although the engineered strain of *C. beijerinckii* (*C. beijerinckii*_mgsA+mgR) expressing a native methylglyoxal synthase (*mgsA*) and a glyoxal/methylglyoxal reductase (*mgR*) from *Clostridium pasteurianum* did not produce 1,2-PDO, it exhibited 35% more growth and produced at least, 85% more butanol than the control strains (*C. beijerinckii*_p459: empty plasmid control and *C. beijerinckii*_mgsA: expressing the native *mgsA*; 16).

In this study, we sought to understand the molecular underpinnings of enhanced butanol production by *C. beijerinckii*_mgsA+mgR relative to the empty plasmid control strain. Using RNA sequencing, we profiled the global mRNA levels in *C. beijerinckii*_mgsA+mgR grown on 50 g/L lactose in comparison to *C. beijerinckii*_p459. Transcriptomic results showed that *de novo* NAD biosynthesis via L-aspartate was strongly downregulated in *C. beijerinckii*_mgsA+mgR. Hence, cultures of *C. beijerinckii*_mgsA+mgR were supplemented with aspartic acid (2 g/L), leading reduced growth and butanol production. We discuss the implications of observed mRNA profiles in this strain, in relation to the observed solvent profiles during lactose catabolism with and without aspartate supplementation.

## MATERIALS AND METHODS

### RNA library construction and sequence analysis

We previously reported the engineering of the strains of *C. beijerinckii* used in this study (16). To isolate total RNA, *C. beijerinckii*_p459 and *C. beijerinckii*_mgsA+mgR were grown in P2 medium as previously described (11), with a slight modification. In place of 60 g/L glucose, the P2 medium contained 50 g/L lactose as carbon source. All cultures were supplemented with erythromycin (15 μg/mL). After the cultures reached an OD_600_ of ∼1.0, triplicate cell samples (10 mL) were collected for RNA isolation. The cells were centrifuged for 10 min at 4000xg. Afterwards, total RNA was extracted with Trizol according to a previously reported method (11). DNA contamination was removed by DNase I treatment (NEB, USA). The quality and quantity of RNA samples were assessed using NanoDrop One UV-Vis Spectrophotometer (ThermoFisher Scientific, Waltham, MA, USA). Ribosomal RNA (rRNA) depletion, library preparation, mRNA sequencing and initial data processing were performed commercially by SeqCoast Genomics (Portsmouth, NH, USA). In brief, rRNAs were removed with the Ribo-Zero Plus Microbiome and then, unique dual index libraries were generated. Sequencing was performed on the Illumina NextSeq2000 platform using a 300 cycle flow cell kit to produce 2×150bp paired reads. Read de-multiplexing, read trimming, and run analytics were performed using DRAGEN v3.10.12, an on-board analysis software on the NextSeq2000. Sequencing quality control was assessed by FastqQC metrics. The reads trimming tool Trimmomatic (version 0.39; 17) was used to remove adaptors and low-quality bases and reads, and Kallisto version 2.1.2 (18) used to generate count tables and normalized expression levels for each of the genes from the reference genome (*C. beijerinckii* NCIMB 8052). DeSeq2 version 2.23.0 (19) was employed to estimate differentially expressed genes and compare gene expression levels in *C. beijerinckii*_mgsA+mgR versus *C. beijerinckii*_p459. Volcano plot was generated using EnhancedVolcano version 2.3, package R (20). BAKTA version 1.5.1 (21) was used for functional annotation of genes. The mRNAs with fold-changes >2 (Log_10_) and a *p*-value <0.05 were considered to be differentially expressed.

### Real-time quantitative PCR

For the real-time quantitative PCR (RT-qPCR) assay, RNA was isolated as described above from triplicate cultures of *C. beijerinckii*_mgsA+mgR and *C. beijerinckii*_p459 grown on lactose and supplemented with erythromycin (15 μg/mL). Using the RNA as template, cDNA was synthesized using the iScript™ cDNA synthesis kit (BioRad, Inc. Hercules, CA, USA). RT-qPCR was performed as per the method described previously (11) using gene specific primers (Table 1) and iTaq Universal SYBR Green Supermix (Bio-Rad, Hercules, CA, USA) in a CFX Connect RT-qPCR System (Bio-Rad, Hercules, CA, USA). Relative expression was normalized relative to those obtained for the *rpoD* housekeeping gene (Cbei_0853) of *C. beijerinckii* and the fold-change of selected genes was calculated using the 2^−ΔΔCt^ method. The data presented is average values of three biological replicates ± standard deviation.

**Table 1.**
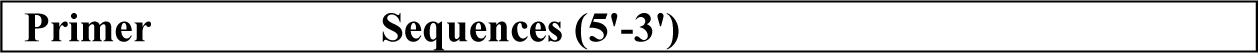
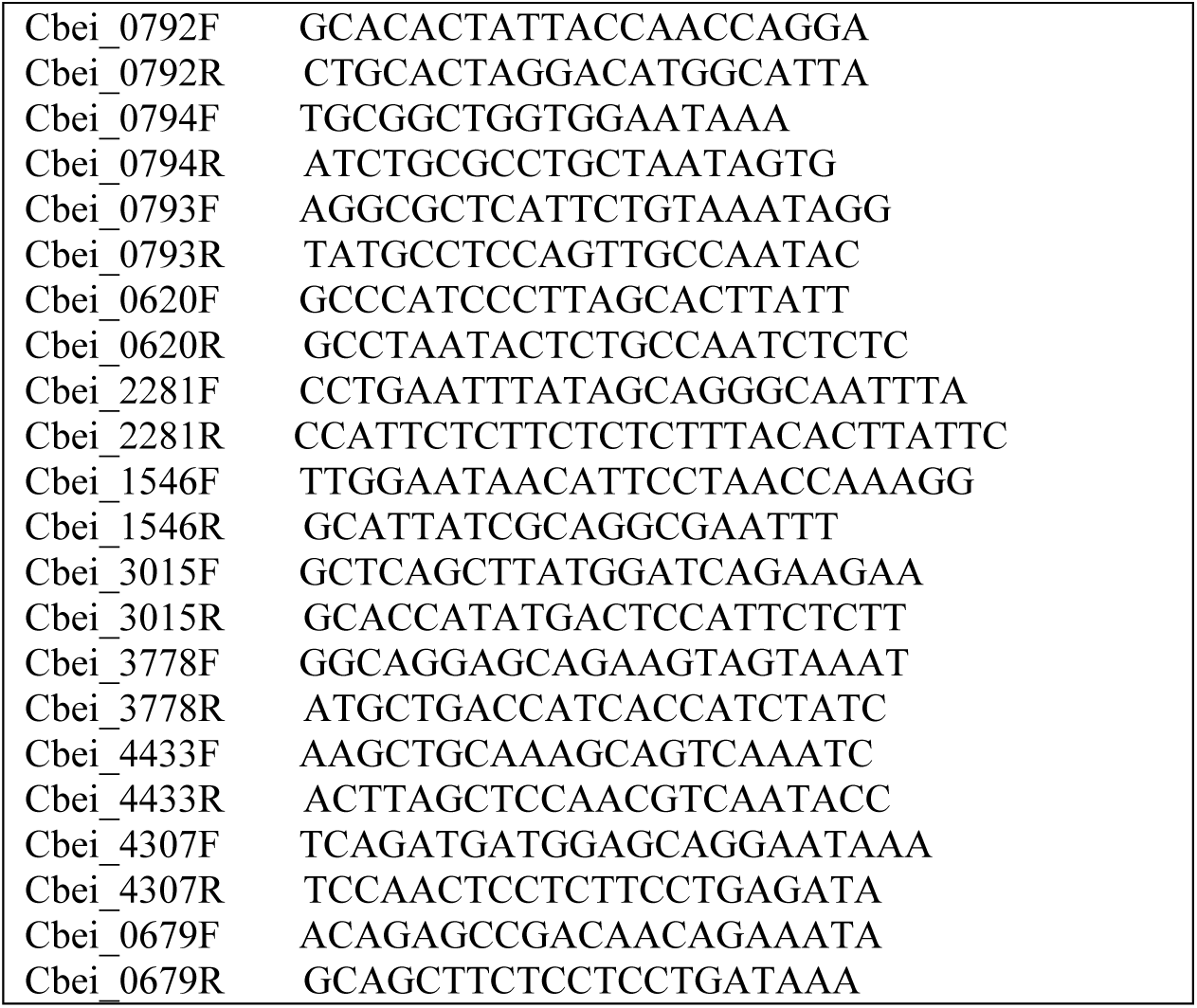
List of primers used for RT-qPCR analysis.

### Aspartic acid supplementation of the fermentation medium

*C. beijerinckii*_mgsA+mgR and *C. beijerinckii*_p459 were grown in lactose-based P2 medium as described earlier. Fermentation on lactose was conducted for 84 h. All cultures were supplemented with aspartic acid (2 g/L) and erythromycin (15 μg/mL). Samples were collected every 12 h and analyzed for the concentrations of acetone, butanol, ethanol, acetic acid, and butyric acid as previously reported (11) using gas chromatography. Concomitantly, samples were analyzed for pH and OD_600_ according to previously reported protocols (11).

### Confocal microscopy

A Nikon Eclipse Ti series microscope (Nikon, Melville, NY, USA) with a 100x oil immersion objective lens was used to capture images on a 1% agarose pad. Two microliters of the culture was placed on the agarose pad and allowed to dry for ∼5 min. Afterwards, the pads were covered with glass slide and then, viewed with oil immersion objective.

## RESULTS

### Comparative transcriptomic profiles of *C. beijerinckii*_mgsA+mgR relative to *C. beijerinckii*_p459

A total of 300 and 433 genes were up- and down-regulated, respectively in *C. beijerinckii*_mgsA+mgR relative to *C. beijerinckii*_p459 (**Table 2**). **Figure 1** is a summary of the distribution of differentially expressed genes in *C. beijerinckii*_mgsA+mgR relative to *C. beijerinckii*_p459. Some of the most strikingly upregulated genes in *C. beijerinckii*_mgsA+mgR include Cbei_3015, Cbei_3778, Cbei_0533, *rpiB*, *crt*, *aroH*, *feoA*, *feoB*, *iolB*, *iolC*, *iolD*, *iolE*, *iolJ*, *CheA*, *fliI*, *CheA*, and *CheB* (**Fig 1**). These genes are involved in electron transfer (e.g., Cbei_3015: flavodoxin), iron uptake, carbohydrate, protein, lipid, or nucleotide metabolism, glycosylation (e.g., Cbei_3778: glycosyl transferase; 22), and DNA recombination (Cbei_0533, recombinase).

**Fig 1.**
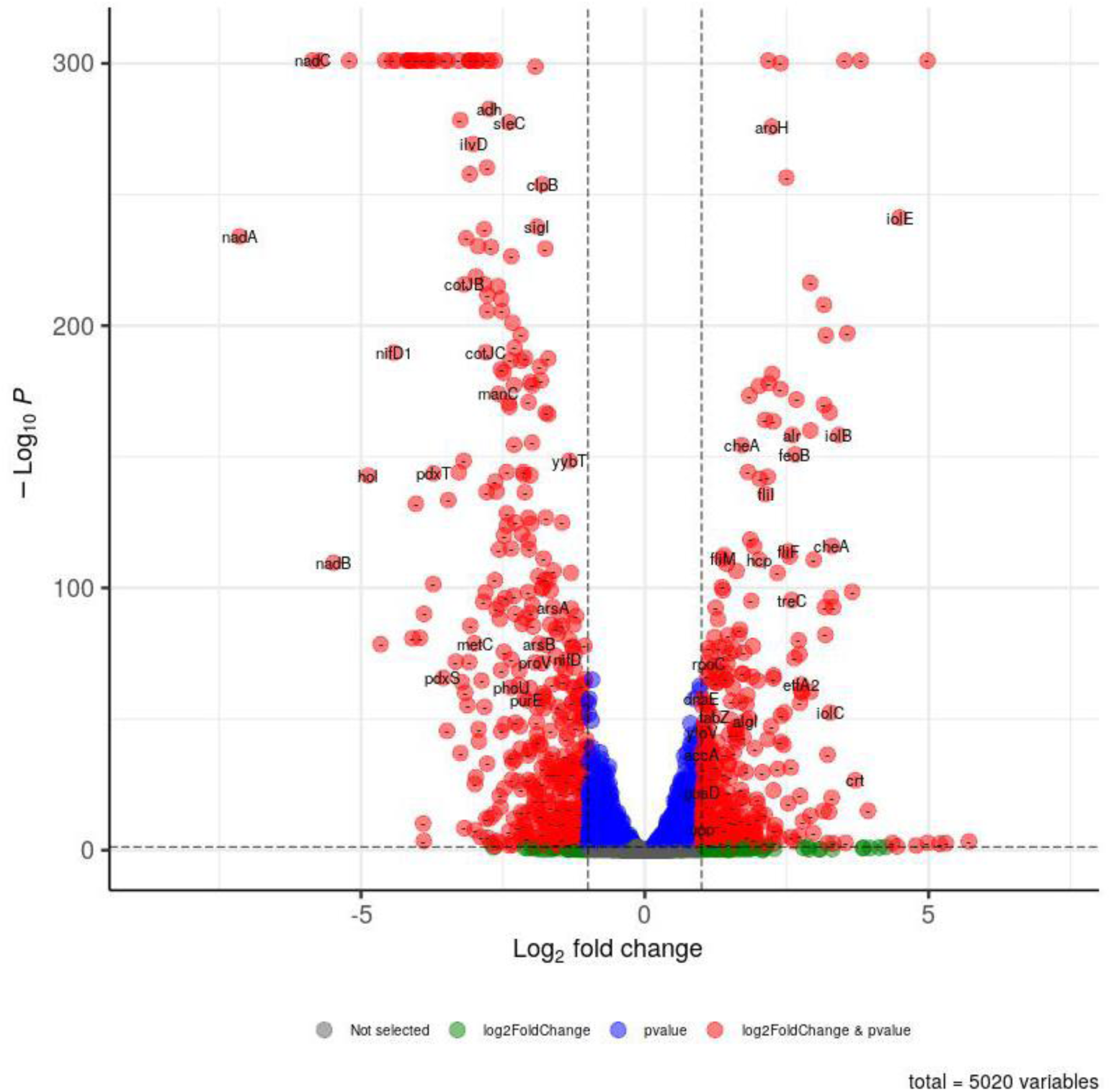
Volcano plot of the differentially expressed genes in *C. beijerinckii*_mgsA+mgR relative to *C. beijerinckii*_p459.

**Table 2.** A summary of the up- or down-regulated genes in *C. beijerinckii*_mgsA+mgR versus *C. beijerinckii*_p459.

|  | Upregulated genes |  | Downregulated genes |  |
| --- | --- | --- | --- | --- |
|  | No. of genes | Percentage (%) | No. of genes | Percentage (%) |
| Total genes | 300 | 100 | 434 | 100 |
| Coenzyme metabolism | 5 | 1.70 | 7 | 1.61 |
| Energy production and conservation | 22 | 7.33 | 29 | 6.68 |
| Intracellular trafficking and secretion | 3 | 1.00 | 2 | 0.46 |
| Secondary structure | 0 | 0.00 | 5 | 1.15 |
| Transcription, replication and repair | 22 | 7.33 | 42 | 9.70 |
| Translation | 6 | 2.00 | 6 | 1.40 |
| Post-translational modification/protein turnover/chaperone functions | 0 | 0.00 | 6 | 1.40 |
| Cell motility and signal transduction | 52 | 17.33 | 15 | 3.50 |
| Nutrient/nucleotide transport and metabolism | 82 | 27.33 | 99 | 22.90 |
| Cell wall/membrane/envelop biogenesis | 10 | 3.33 | 24 | 5.53 |
| Lipid biosynthesis & metabolism | 14 | 4.70 | 10 | 2.30 |
| Cell cycle control and mitosis | 10 | 3.33 | 15 | 3.50 |
| Stress response | 2 | 0.70 | 8 | 1.84 |
| Unknown functions | 73 | 24.33 | 166 | 38.25 |

The highest number of genes (82) upregulated in *C. beijerinckii*_mgsA+mgR relative to *C. beijerinckii*_p459 were genes involved in nutrient/nucleotide transport and metabolism (27.33%). The second group of highly upregulated genes (73) encode proteins of unknown functions (24.33%), followed by genes that encode proteins involved in motility and signal transduction (52/17.33%), transcription, replication and DNA repair (22/7.33%), energy production and conservation (21/7.00%), lipid biosynthesis and metabolism (14/4.70%), and cell wall/membrane biogenesis (10/3.33%). Others include cell cycle control and mitosis (10/3.33%), translation (6/2.00%), coenzyme metabolism (5/1.70%), intracellular trafficking and secretion (3/1.00%), and stress response (2/0.70%; **Table 2**). Genes encoding proteins of unknown function were the most downregulated group of genes in *C. beijerinckii*_mgsA+mgR relative to *C. beijerinckii*_p459 (166/38.34%), followed by genes involved in nutrient/nucleotide transport and metabolism (96/22.20%), transcription, replication and repair (42/9.70%), cell wall/membrane/envelop biogenesis (28/6.50%), energy production and conservation (28/6.50%), cell motility and signal transduction (15/3.50%) and cell cycle control and mitosis (14/3.23%;**Table 2**). Except where stated otherwise, gene expression/mRNA fold changes are presented in log_2_.

#### Differentially expressed nutrient/nucleotide transport and coenzyme metabolism genes

The upregulated nutrient/nucleotide transport and metabolism genes are largely involved in carbohydrate, amino acid, nucleotide, and iron import and metabolism, and peptide processing (**Table S1**). Notably, 17 (∼16.04%) of the upregulated nutrient/nucleotide transport and metabolism genes encode key enzymes/proteins of the phosphotransferase system (PTS) or the iron acquisition machinery (**Table 3**). In fact, 23 of the upregulated nutrient import and metabolism-related genes are involved in carbohydrate import and metabolism (28.04%). Conversely, genes involved in *de novo* biosynthesis of NAD (via L-aspartate) and vitamin B_6_ were downregulated in *C. beijerinckii*_mgsA+mgR relative to *C. beijerinckii*_p459, whereas those encoding proteins involved in the biosynthesis of thiamine and cobalamin (vitamin B_12_) biosynthesis were mostly upregulated (**Tables 4, S1, and S2**). Furthermore, aminodeoxychorismate lyase (also known as 4-amino-4-deoxychorismate lyase; *mltG*), which is involved in folate biosynthesis (**25**) showed 1.20-fold increase in mRNA abundance (**Table S1**). Genes involved in the biosynthesis of coenzyme A, molybdenum cofactor and pantothenate (vitamin B_5_) exhibited irregular patterns of expression. For instance, while *ilvD* (Cbei_1510; dihydroxy-acid dehydratase) involved in the biosynthesis of valine, leucine, isoleucine, vitamin B_5_ and coenzyme A was downregulated 3.03-fold, *coaD* (Cbei_1160; pantothenate-phosphate adenylyltransferase), which also takes part in the biosynthesis of vitamin B_5_ and coenzyme A was upregulated 1.32-fold (**Tables 4, S1, & S2**). The gene *moaC* (Cbei_3792; molybdenum cofactor biosynthesis protein C) showed 1.97-fold reduced mRNA abundance in *C. beijerinckii*_mgsA+mgR, whereas *moaA* (Cbei_1989; molybdenum cofactor biosynthesis protein A) was up-regulated 1.24-fold in the same strain (**Tables 4, S1, & S2**).

**Table 3.**
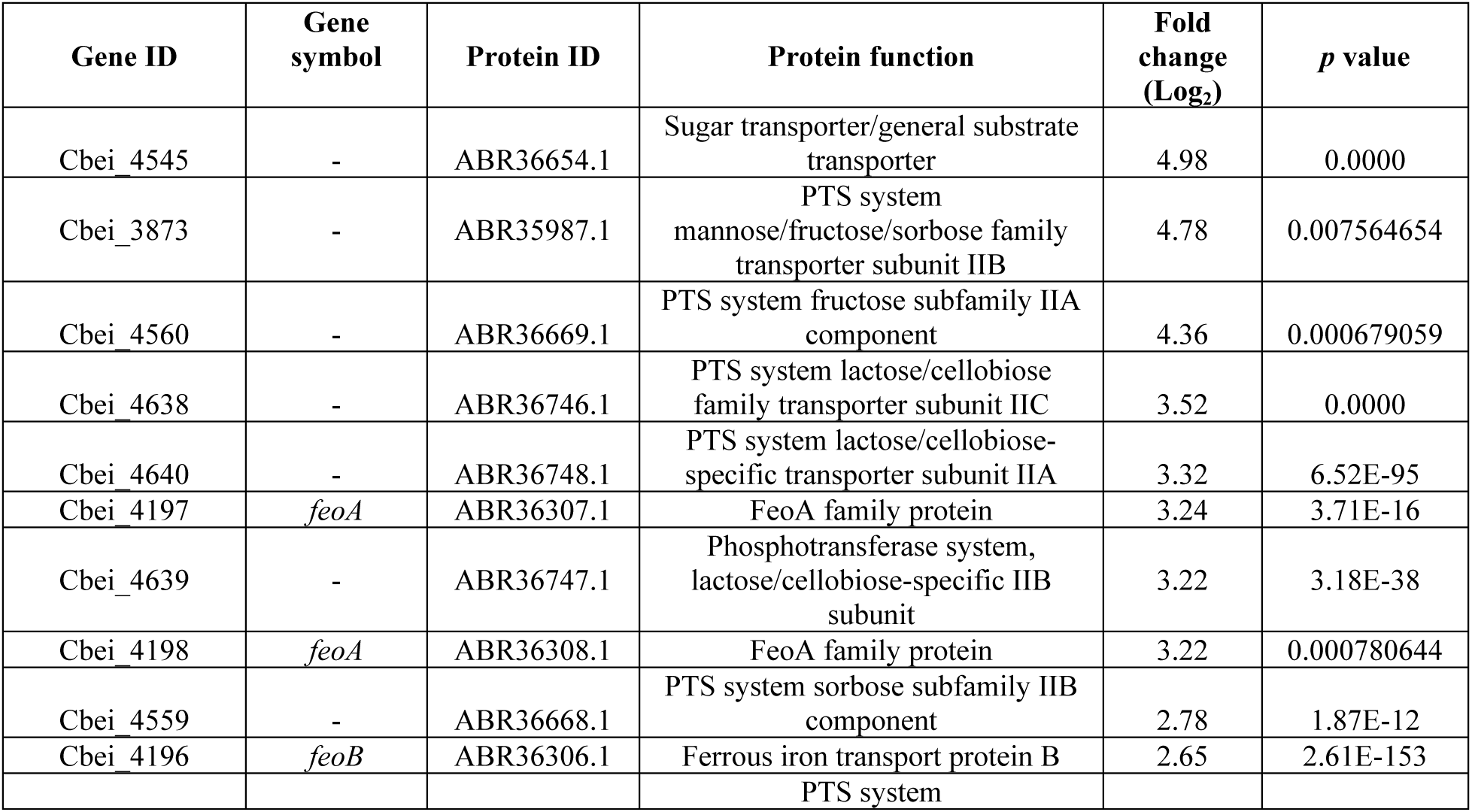
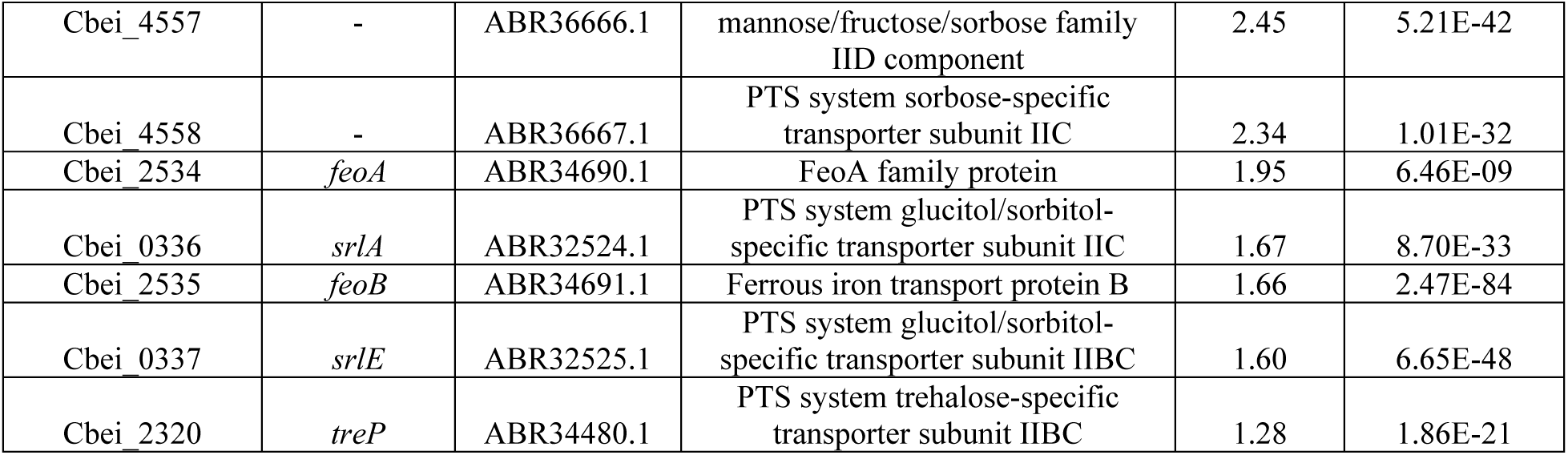
Genes of the phosphotransferase system or iron acquisition machinery upregulated in *C. beijerinckii*_mgsA+mgR relative to *C. beijerinckii*_p459.

**Table 4.**
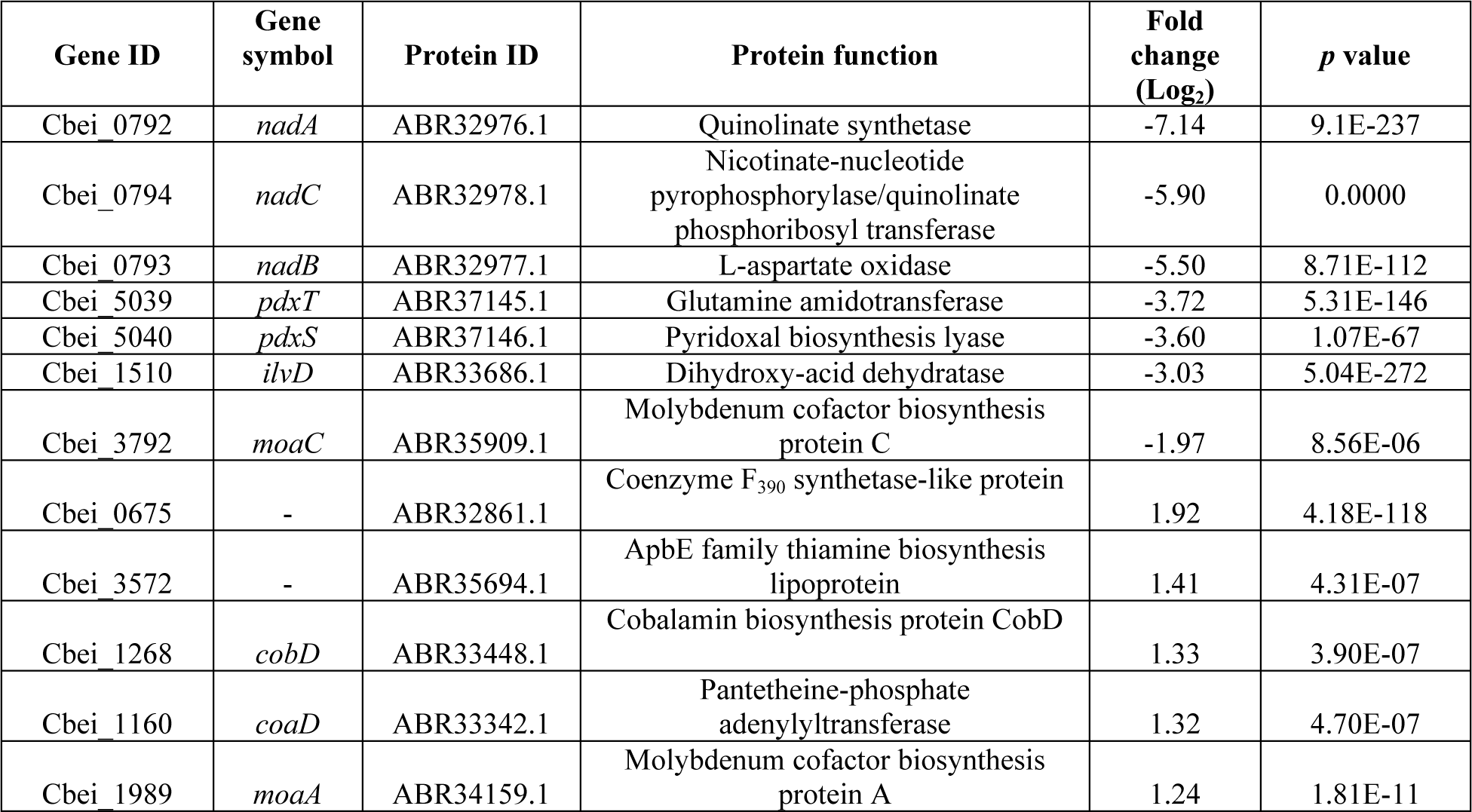
The mRNA levels of genes involved in biosynthesis of coenzymes [NAD, thiamine, cobalamin (thiamin B_12_), molybdenum cofactor, vitamin B_6_, and pantothenate] in *C. beijerinckii*_mgsA+mgR relative to *C. beijerinckii*_p459.

Genes of the myo-inositol catabolic operon (*iolB,C,D,E,J*) were particularly upregulated in *C. beijerinckii*_mgsA+mgR, with fold increases in mRNA abundance ranging from 3.15 – 4.49 (**Table S1**). A significant number of lactose/galactose-specific permeases and general carbohydrate catabolic genes were upregulated in *C. beijerinckii*_mgsA+mgR when compared to *C. beijerinckii*_p459. Notably, a general sugar transporter (Cbei_4545), a monosaccharide-transporting ATPase (*mglC*), a sugar-specific permease (Cbei_3367), and 6-phosphogluconate dehydrogenase (Cbei_2192) were significantly upregulated (**Table S1**). Other sugar transport or catabolic genes upregulated in *C. beijerinckii*_mgsA+mgR were β-glucosidase (Cbei_4681), L-fucose isomerase (Cbei_4681), sucrose-6-phosphate hydrolase (Cbei_3877), transketolase (*dxs_3*), and galactoside O-acetyltransferase (Cbei_4734). Genes associated with biosynthesis of aromatic amino acids (tyrosine, tryptophan and phenylalanine) were upregulated in *C. beijerinckii*_mgsA+mgR. Examples include *aroH* (Phospho-2-dehydro-3-deoxyheptonate aldolase; 2.24-fold), *trpC* (Indole-3-glycerol-phosphate synthase; 1.47-fold), and anthranilate synthase component I (*trpE*), a key enzyme in the indole/tryptophan biosynthesis pathway (23; 1.25-fold; Table S1). Carbamoylphosphate synthase [small (*carA*) and large (*carB*)] subunits were upregulated 1.6- and 1.42-fold, respectively. Carbamoylphosphate synthase is a highly conserved enzyme that catalyzes a step in *de novo* synthesis of arginine and pyrimidines (24).

The mRNA abundance of an amino acid/peptide transporter gene (Cbei_2122) increased 1.90-fold. Conversely, Cbei_0622, *mccA* (Cbei_0630), and *metC* (Cbei_0629), which code for cysteine synthase, cysteine synthase, and cystathionine gamma-synthase involved in cysteine and methionine biosynthesis were downregulated 3.90-, 2.20, and 3.00-fold, respectively (**Table S2**). In addition, *csd* (Cbei_0585) and *csd2* (Cbei_2599), both of which code for cysteine desulfurase were downregulated 2.60- and 1.40-fold, respectively. Likewise, the gene coding for O-acetylhomoserine aminocarboxypropyltransferase (MetY; Cbei_3543) showed 2.25-fold reduced mRNA abundance. MetY is involved in cysteine and methionine metabolism. In addition to downregulation of cysteine and methionine metabolic genes, *asnB* (asparagine synthase) was downregulated 2.33-fold. Similarly, serine--pyruvate transaminase (Cbei_0759), which participates in glycine, serine and threonine metabolism was downregulated 2.30-fold. The gene, *dapB* (dihydrodipicolinate reductase gene) involved in lysine biosynthesis was downregulated 1.66-fold (30), whereas the mRNA levels of *asd* (aspartate-semialdehyde dehydrogenase gene), which plays a crucial role in amino acid and metabolite biosynthesis (32) reduced 1.33-fold. In the same vein, *lysA* (diaminopimelate decarboxylase; DAPDC) was downregulated 1.33-fold. DAPDC catalyzes the last step in bacterial diaminopimelate biosynthesis pathway, which leads to lysine and peptidoglycan biosynthesis in Gram positive bacteria (33). The mRNA levels of other genes that code for lysine biosynthesis enzymes such as *dapA* (dihydrodipicolinate synthase) and *aepX* (cytidyltransferase) reduced 1.30- and 1.27-fold, respectively (**Table S2**).

Pyruvate formate lyase (*pfl*) was upregulated 1.23-fold in *C. beijerinckii*_mgsA+mgR relative to *C. beijerinckii*_p459 (**Table S1**). The protein product of *pfl* (PFL) supplies acetyl-CoA from pyruvate and CoA during anaerobic glycolysis (26). PFL is activated by an activase that belongs to the radical SAM (*S*-adenosylmethionine) protein superfamily (26). Radical SAM proteins generate a radical species via reductive splitting of SAM by deploying an uncommon Fe-S center (26–27). Although genes for 2 radical SAM proteins (Cbei_0628 and Cbei_0037; **Table S2**) were downregulated, genes coding for 3 radical SAM proteins (Cbei_0683, Cbei_0690, and Cbei_3030) were upregulated (**Table S1**). Furthermore, *nrdG* (anaerobic ribonucleoside-triphosphate reductase activating protein; ARTRAP) showed a 1.33-fold increase in mRNA abundance. ARTRAP activates anaerobic ribonucleoside-triphosphate reductase during anaerobic growth by generating an organic-free radical with SAM and reduced flavodoxin as co-substrates to produce 5’-deoxy-adenosine (28).

Genes associated with uptake of purines and pyrimidines were upregulated in *C. beijerinckii*_mgsA+mgR. For example, the mRNAs levels of the ORFs, Cbei_1974 (xanthine/uracil/vitamin C permease) and Cbei_1715 (*pyrP*; uracil-xanthine permease) increased 1.20- and 1.26-fold, respectively (**Table S1**). Furthermore, Cbei_1982 (aldehyde oxidase/xanthine dehydrogenase; molybdopterin-binding subunit) was upregulated 1.20-fold. Conversely, *pbuX*, (uracil-xanthine permease) showed 1.20-fold reduced mRNA abundance. Conversely, the mRNA abundance levels of *purE* (phosphoribosylaminoimidazole carboxylase catalytic subunit) that catalyzes the sixth step in *de novo* purine biosynthesis reduced 2.10-fold (**Table S2**). Concomitantly, *purL* and *purH* (phosphoribosylformylglycinamidine synthase and 5-aminoimidazole-4-carboxamide ribonucleotide transformylase, respectively) that catalyze the fourth and the last two steps of *de novo* purine biosynthesis were downregulated 1.27- and 1.22-fold, respectively.

A considerable number of iron-dependent proteins were downregulated in *C. beijerinckii*_mgsA+mgR, compared to *C. beijerinckii*_p459. Although not to same degree, this was accompanied by considerable upregulation of some iron-dependent proteins, one of which includes the gene for a hemerythrin-like metal-binding protein (Cbei_2165), which was upregulated 1.69-fold. Among an assortment of functions, hemerythrin-like proteins participate in iron homeostasis (29). Oddly, 5 genes that code for ferrous iron uptake proteins (3 *feoA*s and 2 *feoB*s) were strongly upregulated, with increases in mRNA fold changes ranging from 1.66- to 3.25-fold (**Table S1**). Similarly, two copies of *phoU* (Cbei_1131 and Cbei_1132; **Table S2**) were significantly downregulated in *C. beijerinckii*_mgsA+mgR. The protein product of *phoU* (PhoU) is a negative regulator of Pho regulon genes under phosphate-replete conditions (**Table S2**). Concomitantly, *ppk1* (polyphosphate kinase) was downregulated 2.61-fold. Molybdenum ABC transporter (*modA*) exhibited 1.97-fold reduced mRNA abundance. The gene for an amine oxidase (Cbei_2598), which catalyzes the conversion of primary amines to aldehydes (31) was downregulated 1.33-fold, whereas *amt* (ammonium transporter) was downregulated 1.20-fold.

#### Expression patterns of cell motility and signal transduction genes

Genes that code for proteins involved in cell motility and signal transduction accounted for the third most upregulated category of genes (**Table S1**). Among the upregulated cell motility and signal transduction genes, genes which encode proteins associated with biosynthesis and assembly of flagellum components and functioning of the flagellum accounted for 25.00%. Similarly, 25.00% of the upregulated cell motility and signal transduction genes code for methyltransferases and related proteins involved in the reception, processing, and transmission of signals from chemoreceptors to the flagella motors. Histidine kinases and methyl-accepting chemotaxis sensory transducers (MCPs) accounted for 17.00% each, of the upregulated cell motility and signal transduction genes. Conversely, only 15 of the downregulated genes (3.50%) are associated with cell motility and signal transduction, out of which 5 genes code for MCPs (**Table S2**). Notably, 9 MCPs were upregulated in *C. beijerinckii*_mgsA+mgR in comparison to *C. beijerinckii*_p459. Two diguanylate cyclase/phosphodiesterase genes were downregulated in *C. beijerinckii*_mgsA+mgR relative to *C. beijerinckii*_p459, whereas one diguanylate cyclase with PAS/PAC sensor was upregulated in the same strain.

#### The mRNA profiles of energy production and conservation genes

A similar number of genes involved in energy production and conservation were up-(22/7.33%) and down-regulated (28/6.50%) in *C. beijerinckii*_mgsA+mgR when compared to *C. beijerinckii*_p459 (**Table 2**). However, electron transfer FAD-dependent enzymes/proteins were predominantly upregulated in *C. beijerinckii*_mgsA+mgR (12/54.60%; **Table S1**), including flavodoxin, which showed the greatest increase (5.71-fold) in mRNA abundance relative to *C. beijerinckii*_p459. Notably, 23.81% of the upregulated genes involved in energy production and conservation encode iron-dependent proteins [e.g., iron-dependent alcohol dehydrogenases (Cbei_4552 and Cbei_2194), Fe-S (4Fe-4S) cluster carbon-monoxide dehydrogenase (Cbei_5054), Fe(Cys)4 rubrerythrin (Cbei_1416), and Fe-only hydrogenase (Cbei_4110)]. Additionally, 4 NAD(H)-dependent iron-containing proteins (including D-isomer specific 2-hydroxyacid dehydrogenase - Cbei_2193; NADH-ubiquinone oxidoreductase subunit 6 - Cbei_2990; aldehyde dehydrogenase - Cbei_0674, and NADH dehydrogenase - Cbei_4112) were also upregulated. Conversely, 13 genes that encode iron-containing/dependent proteins (46.43%) involved in energy production and conservation, including a rubredoxin-type Fe(Cys)4 protein (Cbei_1012), an iron-containing alcohol dehydrogenase (Cbei_2181), a nitrogenase molybdenum-iron protein (alpha chain; Cbei_2002), a dinitrogenase iron-molybdenum cofactor biosynthesis protein (Cbei_0627), one nitroreductase (Cbei_3643), and 2 oxidoreductases/nitrogenases (Cbei_0620/*nifD1* and Cbei_0621) were downregulated in *C. beijerinckii*_mgsA+mgR (**Table S2**). Markedly, 7 of the 14 downregulated genes that code for iron-containing/dependent proteins associated with energy production and conservation are ‘iron-intensive’ Fe-S (4Fe-4S/2Fe-2S) cluster proteins (**Table S2**). For instance, three genes (Cbei_0795, Cbei_2128 and Cbei_3544, and Cbei_4835) that code for (4Fe-4S) ferredoxin iron-sulfur binding domain protein, ferredoxin-NADP^+^ reductase subunit alpha, and 2 4Fe-4S ferredoxins, respectively, which orchestrate electron transfer, particularly to hydrogenases were significantly downregulated.

#### The expression levels of genes involved in transcription, replication, repair and translation

As shown in **Table 2**, the mRNA levels of genes involved in transcription, replication, repair and translation (TRRT) were the 4^th^ and 3^rd^ most up- and down-regulated categories, respectively, during the growth of *C. beijerinckii*_mgsA+mgR on lactose. Although TRRT genes accounted for similar percentages of the up- (9.33%) and down-regulated (11.10%) genes in *C. beijerinckii*_mgsA+mgR, the actual number of downregulated TRRT genes (48) was 71.42% higher than those that were upregulated (28). Conspicuously, 3 periplasmic binding protein/LacI transcriptional regulators (Cbei_4434, Cbei_2377 and Cbei_0334) involved in lactose metabolism were strongly upregulated (**Table S1**). Accessory regulator proteins (Cbei_4528 and Cbei_0269) that promote the expression of exoproteins and repression of some cell wall-associated proteins (**35**), and a recombinase (Cbei_0533) were upregulated (**Table S1**). Other upregulated TRRT genes in *C. beijerinckii*_mgsA+mgR include those involved in transcriptional regulation of signal transduction (ECF subfamily RNA polymerase sigma-24 factor; Cbei_1302), glucitol (sorbitol) operon activation (glucitol operon activator; Cbei_0335), β-glucoside utilization (*bgl*; transcriptional antiterminator BglG), protein acetylation and stress response (GCN5 N-acetyltransferase; Cbei_2936). Gene for the DeoR family transcriptional regulator (Cbei_0466), a repressor of sugar metabolism (**36**), sporulation stage II, protein E (*spoIIE*; Cbei_0097), MarR family transcriptional regulator (Cbei_0302), and nuclease SbcCD subunit D (*sbcD*; Cbei_0980) that facilitate DNA transcription exhibited 1 increases in mRNA abundance (**Table S1**).

Eight xenobiotic response element family transcriptional regulators were notably downregulated in *C. beijerinckii*_mgsA+mgR (**Table S2**). Further, the mRNA levels of the iron-dependent transcription repressor (*ideR*; Cbei_4069), which regulates intracellular free iron concentration (**37**) decreased 3.20-fold. In addition, genes involved in transcriptional regulation of sporulation (*abrB*; Cbei_2219; 2.60-fold), signal transduction (RNA polymerase, sigma-24 subunit, ECF subfamily; Cbei_3743 and a serine/threonine kinase related protein; Cbei_1424), carbon metabolism (*rpiR*; Cbei_1598; 1.90-fold), intracellular trafficking, organelle biogenesis, cytoskeleton regulation, DNA replication and intracellular motility (**38**; AAA ATPase; Cbei_3732) were downregulated. The genes *sigI* (RNA polymerase sigma factor 70; Cbei_1826), *marR* family transcriptional regulator (Cbei_3301; 1.85-fold), *ecf* subfamily RNA polymerase sigma-24 factor (Cbei_2303), a two component transcriptional regulator (Cbei_2951), *lysR* family transcriptional regulator (Cbei_3631; 1.55-fold), and *hutP* (Cbei_1078) also exhibited varying degrees of mRNA downregulation. These genes are involved in transcriptional regulation of metabolism, stress response, signal transduction, DNA replication, quorum sensing and motility.

Three GCN5-related N-acetyltransferases (Cbei_3937, Cbei_3291, and Cbei_3844) showed considerable reductions in relative mRNA abundance (**Table S2**). Furthermore, 4 *tetR* family transcriptional regulators (Cbei_3644, 1Cbei_2866, Cbei_1253, and Cbei_2924) that code for the AcrR family transcriptional regulator exhibited decreased mRNA abundance. The products of these genes are associated with the regulation of cellular metabolism, biofilm formation and efflux. Also downregulated was the gene for LuxR family transcriptional regulator (Cbei_3706), which controls the expression of a wide range of genes involved in biofilm formation, motility, and nodulation (**39**). In addition, the genes for a MerR family transcriptional regulator (associated with Hg resistance; Cbei_2721), BadM/Rrf2 family transcriptional regulator (Cbei_2221), *ctsR* (Cbei_0120), a master repressor of stress response gene expression (40), and 2 copies of glucose-inhibited division protein A genes (*gidA*), involved in growth and stress response were downregulated.

### The mRNA levels of genes involved cell wall/membrane/envelop biogenesis

Genes involved in the biosynthesis of cell wall and membrane components were more predominantly downregulated, with a few specific genes involved in peptidoglycan biosynthesis exhibiting greater mRNA abundance. Notably, 4 Glycosyl transferase (GT) genes involved in biosynthesis of cell wall carbohydrates and glycoconjugates (41–42) were upregulated, while 5 GTs were downregulated. Also significantly upregulated were *alr* (alanine racemase; Cbei_4113), 2 copies of membrane bound O-acyl transferase (MBOAT) family protein (*algI*; Cbei_1088 and *dltB*; Cbei_4331), D-alanine--D-alanine ligase (*ddl*; Cbei_0581), and undecaprenyl-phosphate galactose phosphotransferase (Cbei_4575). These genes are involved in biosynthesis of different components of the bacterial cell wall or membrane architectures and their ancillary structures including D-alanine (D-Ala; 43), triacylglycerol, d-alanyl–d-alanine dipeptide (44), and the precursor of O-polysaccharide. Concomitantly, a wide range of genes involved in cell wall/membrane remodeling and biogenesis, capsule formation and growth were downregulated. Notable examples include UDP-glucose 6-dehydrogenase (*wbpA1*; Cbei_4708), mannose-1-phosphate guanylyltransferase (GDP) (*manC*; Cbei_0318), gene for NLP/P60 protein (Cbei_1242), prolipoprotein diacylglyceryl transferase (Cbei_2970), and UDP-N-acetylglucosamine 1-carboxyvinyltransferase (*murA*; Cbei_0421). These genes are largely associated with biosynthesis of capsular polysaccharide. Other downregulated genes associated with cell wall/membrane/capsular structures include glucosamine--fructose-6-phosphate aminotransferase (*glmS*; Cbei_0246) and genes of the L-rhamnose—an important component of cell wall lipopolysaccharide—biosynthesis operon [*rfbA* (Cbei_2602), *rfbB* (Cbei_2578), *rfbC* (Cbei_2601), and *rfbD* (Cbei_2579); **Table S2**)].

### Expression profiles of genes associated with lipid/fatty acid metabolism and biosynthesis

Similar numbers of genes associated with lipid/fatty acid metabolism and biosynthesis were up- (14) and down-regulated (10) in lactose-grown *C. beijerinckii*_mgsA+mgR relative to *C. beijerinckii*_p459 (**Tables S1 & S2**). However, a greater number of the upregulated genes are centrally involved lipid metabolism and fatty acid biosynthesis (**Table S1**). For example, 4’-phosphopantetheinyl transferase (Cbei_0687), acyl-ACP thioesterase (Cbei_0691), oleoyl-(acyl-carrier-protein) hydrolase (Cbei_0248), and thioesterase (Cbei_0681), which are involved in various stages of fatty acid biosynthesis including control of the profile lengths of fatty acids produced by type II fatty acid synthases (45–47) were significantly upregulated in *C. beijerinckii*_mgsA+mgR relative to *C. beijerinckii*_p459. Additionally, beta-ketoacyl synthase-like protein (*fabF*; Cbei_1072), beta-hydroxyacyl-(acyl-carrier-protein) dehydratase (*fabZ*; Cbei_1074), 3-oxoacyl-(acyl-carrier-protein) reductase (*fabG*; Cbei_1071), the three of which are involved in the elongation of fatty acid chains (48) were upregulated. Concomitantly, enoyl-CoA hydratase/isomerase (*crt*; Cbei_4544), acyl-CoA dehydrogenase (Cbei_4542), short-chain dehydrogenase/reductase (*sdr*; Cbei_0340), and lysophospholipase (Cbei_1305) associated with fatty acid/lipid metabolism were upregulated in *C. beijerinckii*_mgsA+mgR, when compared to *C. beijerinckii*_p459. Two alpha/beta fold family hydrolases (Cbei_3932 and Cbei_3997), 3-oxoacid CoA-transferase subunits A and B (*atoD*; Cbei_2654 and *ctfB1*; Cbei_2653), and acetyl-CoA acetyltransferase (Cbei_3630) that take part in fatty acid metabolism were down-regulated.

### Quantitative RT-PCR

A total of 10 differentially expressed genes were studied by reverse transcriptase quantitative PCR (RT-qPCR). As observed with RNA sequencing, the RT-qPCR results showed that Cbei_0792 (quinolinate synthetase - *nadA*), Cbei_0794 (nicotinate-nucleotide pyrophosphorylase/quinolinate phosphoribosyl transferase - *nadC*), Cbei_0793 (L-aspartate oxidase - *nadB*), Cbei_0620 (Oxidoreductase/nitrogenase - *nifD1*), Cbei_2281 (oxidoreductase FAD/NAD(P)-binding domain protein), and Cbei_1546 (hypothetical protein) were downregulated (**Table 5**). On the other hand, Cbei_3015 (flavodoxin), Cbei_4307 (CheA signal transduction histidine kinase - *cheA*), Cbei_3778 (glycosyl transferase family protein), and the ABC transporters (Cbei_4433 – *mglA* and Cbei_0679 – *yfmM*) were significantly upregulated.

**Table 5.** qPCR verification of differentially expressed genes in *C. beijerinckii*_mgsA+mgR relative to *C. beijerinckii*_p459.

| Gene ID | Gene name | Fold change (Log<sub>2</sub>) |  |
| --- | --- | --- | --- |
|  |  | RNA sequencing<br>(*p* value) | qRT-PCR<br>(SD) |
| Cbei_0792 | Quinolinate synthetase (*nadA*) | -7.14 ± 9.1E-237 | -9.39 ± 0.00023874 |
| Cbei_0794 | Nicotinate-nucleotide<br>pyrophosphorylase/quinolinate phosphoribosyl<br>transferase (*nadC*) | -5.90 ± 0.00000 | -6.68 ± 0.00281 |
| Cbei_0793 | L-aspartate oxidase (*nadB*) | -5.50 ± 8.71E-112 | -9.60 ± 0.00024 |
| Cbei_0620 | Oxidoreductase/nitrogenase (*nifD1*) | -4.42 ± 2.82E-192 | -1.40 ± 0.05985 |
| Cbei_2281 | Oxidoreductase FAD/NAD(P)-binding domain<br>protein | -1.91 ± 0.23780 | -0.50 ± 0.39370 |
| Cbei_1546 | Hypothetical protein | -0.999 ± 2.12E-35 | -0.96 ± 0.43073 |
| Cbei_3015 | Flavodoxin | 5.71 ± 0.00016 | 6.50 ± 0.30834 |
| Cbei_4307 | CheA signal transduction histidine kinase (*cheA*) | 1.71 ± 9.07E-157 | 4.16 ± 0.15990 |
| Cbei_3778 | Glycosyl transferase family protein | $5.19 \pm 0.00140$ | $4.08 \pm 0.62201$ |
| Cbei_4433 | ABC transporter related (*mglA*) | $1.90 \pm 6.96\text{E-}80$ | $2.33 \pm 0.24330$ |
| Cbei_0679 | ABC transporter (*yfmM*) | $1.12 \pm 1.31\text{E-}55$ | $1.10 \pm 0.26880$ |

SD – standard deviation ( $n = 3$ )

### Effect of aspartic acid supplementation on the fermentation profile of *C. beijerinckii*_mgsA+mgR

We have previously shown that *C. beijerinckii*_mgsA+mgR exhibits superior growth, butanol and total solvent (acetone-butanol-ethanol – ABE) profiles when compared to the control strain (*C. beijerinckii*_p459; 16). Following strong downregulation of multiple genes involved in *de novo* biosynthesis of NAD in *C. beijerinckii*_mgsA+mgR via the L-aspartate route, we supplemented cultures of this strain with aspartic acid (2 g/L). Remarkably, with aspartate supplementation, OD_600_ of *C. beijerinckii*_mgsA+mgR reduced ∼62%, relative to the cultures of *C. beijerinckii*_p459 (**Fig. 2A**). Concomitantly, butanol and ABE concentrations decreased 434% and 185%, respectively, in cultures of *C. beijerinckii*_mgsA+mgR relative to those of *C. beijerinckii*_p459 (**Fig. 2B & C**). Aspartic acid supplementation impaired acid re-assimilation in *C. beijerinckii*_mgsA+mgR but not in *C. beijerinckii*_p459 (**Fig. 3**). As shown in Fig. 3, addition of aspartic acid to the fermentation medium led to consistently higher concentrations of acetate and butyrate in cultures of *C. beijerinckii*_mgsA+mgR when compared to *C. beijerinckii*_p459 (**Fig. 3 A & B**). Consequently, 19% and 41% more acetate and butyrate, respectively, remained in the cultures of *C. beijerinckii*_mgsA+mgR at the end of fermentation, when compared to the cultures of *C. beijerinckii*_p459. Conversely, in cultures without aspartic acid, both strains exhibited similar patterns of acid re-assimilation, particularly, acetate (**Fig. 3 C & D**). In fact, without aspartic acid supplementation, at the end of fermentation, both *C. beijerinckii*_mgsA+mgR and *C. beijerinckii*_p459 contained 1.39 and 1.35 g/L acetate, respectively, whereas the cultures of *C. beijerinckii*_p459 contained 99% more butyrate (2.13 g/L) than those of *C. beijerinckii*_mgsA+mgR (1.07 g/L).

**Fig 2.**
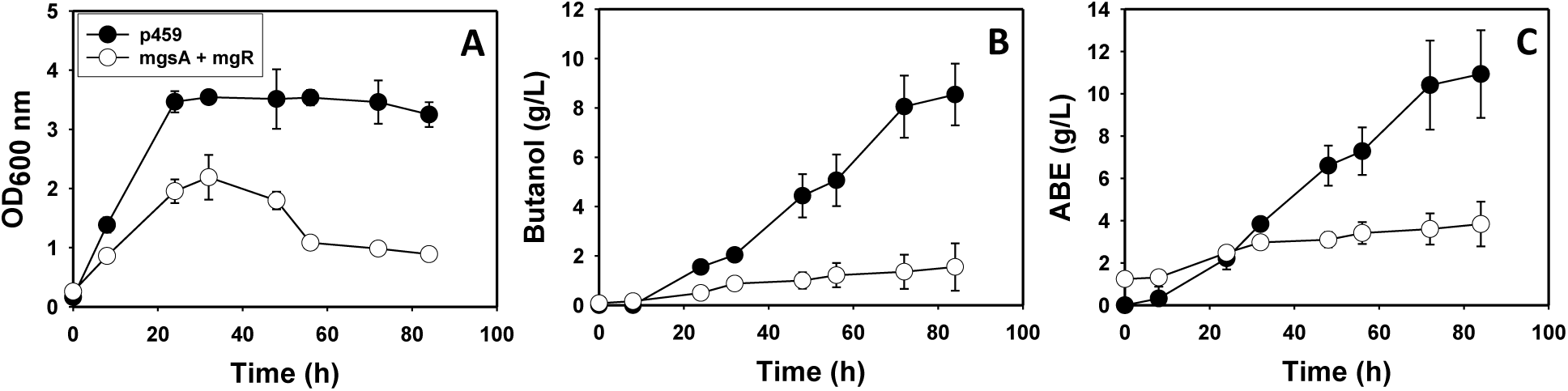
Growth and solvent profiles of *C. beijerinckii*_mgsA+mgR and *C. beijerinckii*_p459. (A) Growth (OD_600 nm_). (B) Butanol concentrations. (C) Total acetone, butanol, and ethanol (ABE) concentrations. (*n* = 3)

**Fig 3.**
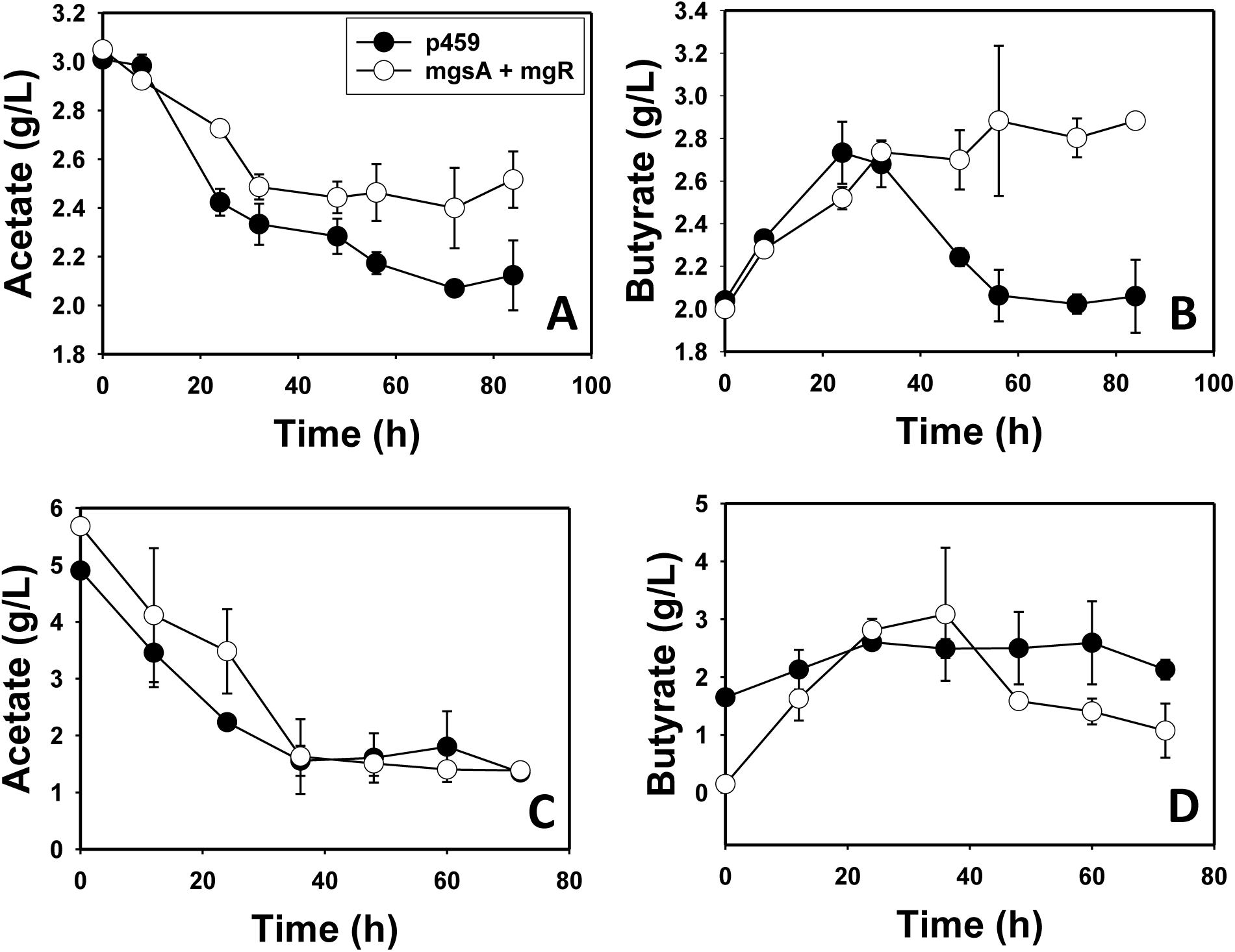
Acetate and butyrate concentrations in aspartate-supplemented and un-supplemented cultures of *C. beijerinckii*_mgsA+mgR and *C. beijerinckii*_p459. (A) Acetate concentrations in aspartate-supplemented cultures. (B) Butyrate concentrations in aspartate-supplemented cultures. (C) Acetate concentrations in aspartate un-supplemented cultures. (D) Butyrate concentrations in aspartate un-supplemented cultures. (n = 3)

### Morphological features of *C. beijerinckii*_mgsA+mgR and *C. beijerinckii*_p459

Examination of *C. beijerinckii*_mgsA+mgR and *C. beijerinckii*_p459 using confocal microscopy revealed that while cells of *C. beijerinckii*_mgsA+mgR appeared predominantly as discrete, individual cells, *C. beijerinckii*_p459 occurred largely in clusters of closely attached cells (**Fig. 4**).

**Fig 4.**
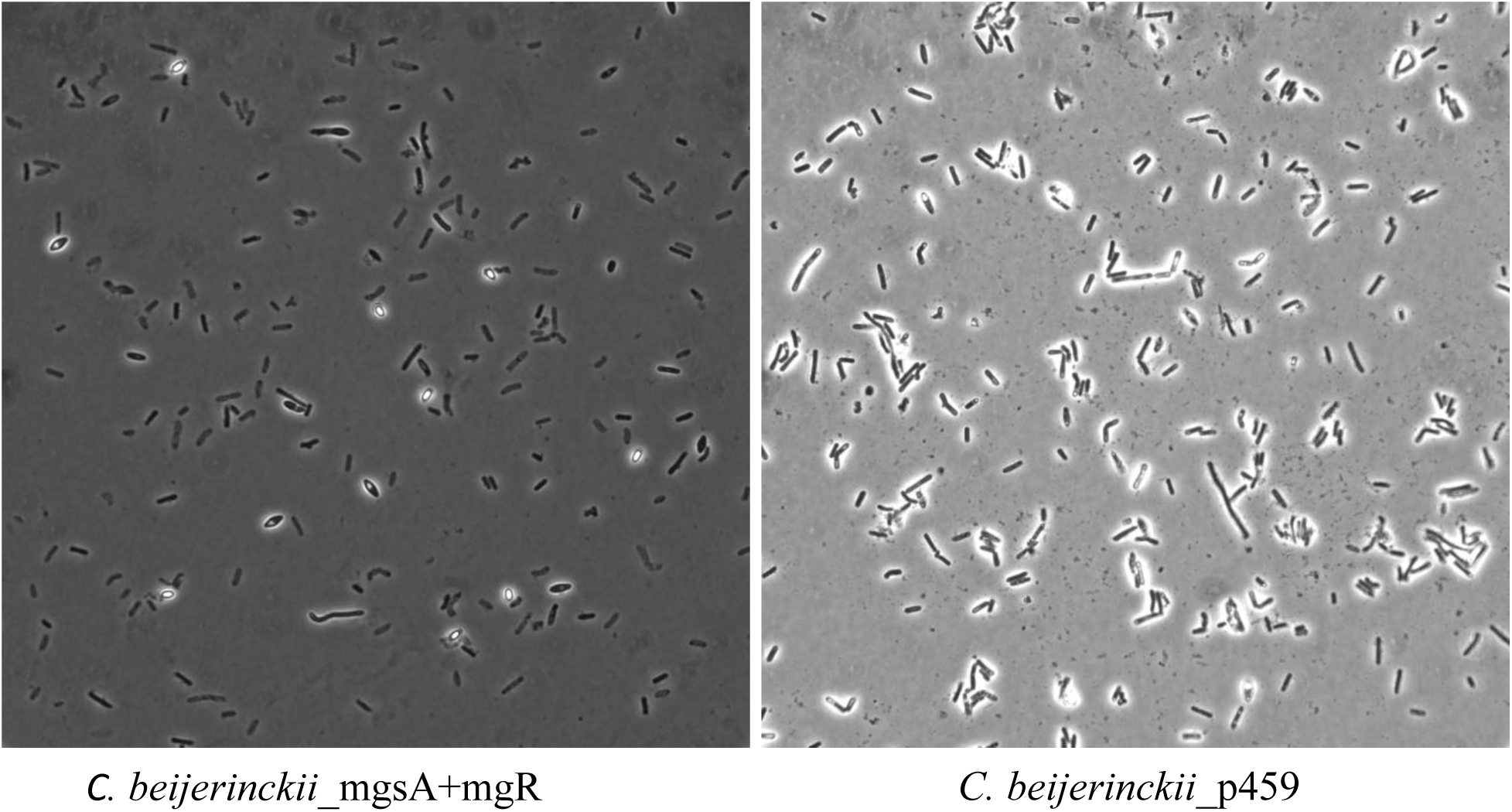
Morphological features and distribution of the cells of *C. beijerinckii*_mgsA+mgR and *C. beijerinckii*_p459 grown on lactose for 24 h.

## DISCUSSION

We previously reported improved growth and butanol production by *C. beijerinckii*_mgsA+mgR (relative to the empty plasmid control strain: *C. beijerinckii*_p459; 16). Here, we present the global transcriptomic profile of *C. beijerinckii*_mgsA+mgR relative to *C. beijerinckii*_p459, in an effort to delineate the underlying molecular basis for enhanced growth and solvent production in lactose-grown *C. beijerinckii*_mgsA+mgR. Broadly, when compared to *C. beijerinckii*_p459, *de novo* biosynthesis of NAD via L-aspartate, vitamin B_6_, purines, and capsular exopolysaccharides were downregulated in *C. beijerinckii*_mgsA+mgR. Similarly, asparagine, cysteine and methionine, glycine, serine, lysine, and threonine metabolism was downregulated in *C. beijerinckii*_mgsA+mgR. Likewise, biosynthesis of valine, leucine, and isoleucine, ammonium uptake and heat shock chaperones were downregulated in *C. beijerinckii*_mgsA+mgR, relative to *C. beijerinckii*_p459. Conversely, phosphate, iron, purine, pyrimidine, and peptide uptake; biosyntheses of thiamine, vitamin B_12_, vitamin B_5_, folate, tyrosine, aromatic amino acids (likely tryptophan), arginine, and pyrimidines were largely upregulated in *C. beijerinckii*_mgsA+mgR. In addition, the mRNA profiles indicate that the two component signal transduction system and cell motility, flavodoxin (and indeed, flavoproteins) and lipid biosynthesis and metabolism were significantly upregulated in *C. beijerinckii*_mgsA+mgR, relative to *C. beijerinckii*_p459.

The transcriptomic repertoire of lactose-grown *C. beijerinckii*_mgsA+mgR suggests that the cells likely experienced lower levels of intracellular ferrous iron, which may have triggered a series of gene expression patterns that led to enhanced growth, lactose utilization and solvent production. Ferrous iron plays a vital role in ABE fermentation. Above a certain threshold (10 mg/L), increasing ferrous iron concentration impairs growth, sugar utilization and solvent production, with concomitant acid accumulation (49). Conversely, lower concentrations of iron favor sugar utilization, growth and ABE production (49). Notably, *C. beijerinckii*_mgsA+mgR consumed 183.33% more lactose than *C. beijerinckii*_p459 (16). It is important to highlight that both *C. beijerinckii*_mgsA+mgR and *C. beijerinckii*_p459 were grown in the same medium containing 10 mg/L FeSO_4_.7H_2_O. However, the expression profiles of iron regulatory and uptake genes in *C. beijerinckii*_mgsA+mgR clearly suggest a concerted cellular effort to increase iron uptake in *C. beijerinckii*_mgsA+mgR. Specifically, the iron-dependent repressor *ideR* (Cbei_4069) was significantly downregulated, when compared to *C. beijerinckii*_p459 (Table S2). IdeR is an iron responsive transcription factor that functions as a gatekeeper to regulate iron uptake (50). In fact, *ideR* was the second most downregulated transcription factor in *C. beijerinckii*_mgsA+mgR. IdeR controls the expression of several genes involved in iron uptake, metabolism, and storage. Significant downregulation of *ideR* is indicative of possibly lower intracellular levels of ferrous iron in *C. beijerinckii*_mgsA+mgR, which might explain increased expression of the iron uptake genes *feoA* (3 copies) and *feoB* (2 copies; Table 3). Other indications of possible ferrous iron crisis in *C. beijerinckii*_mgsA+mgR include widespread downregulation of iron-intensive ([4Fe-4S]/[2Fe-2S]) proteins and upregulation of a hemerythrin-like protein (Table S1). Hemerythrin-like proteins participate in iron homeostasis and have been reported to repair iron centers in *Escherichia coli* (29).

Methylglyoxal (MG), the product of the MgsA-catalyzed reaction is a highly reactive α-oxoaldehyde that exerts severe stress on biological cells, which results in complex formation with proteins via glycation (51–53). Iron has been reported to promote MG-mediated protein damage (52). Therefore, we speculate that the MG product of MgsA likely interacts with ferrous iron and Fe-S proteins, which are abundant in *C. beijerinckii* (54). The resulting iron binding MG-protein complexes likely make free iron less available within the cell (Fig. 5). It is important to note that likely production of MG by MgsA in *C. beijerinckii*_mgsA+mgR did not impair growth. MG was not detected in cultures of *C. beijerinckii*_mgsA+mgR. Possible formation of iron binding MG-protein complexes may account at least in part, for inability to detect MG in cultures of *C. beijerinckii*_mgsA+mgR. Additionally, co-expression of *mgsA* and *mgR* (16) likely resulted in enhanced detoxification of MG by the aldehyde reductase enzyme product of *mgR*. Previously, we observed that a strain of *C. beijerinckii* expressing *mgsA* alone (without the tethered aldehyde reductase in *C. beijerinckii*_mgsA+mgR exhibited diminished growth, and the presence of MG in the culture (16). This assertion is supported by the collective expression profiles of *csd* (cysteine desulfurase) cysteine and methionine biosynthesis and metabolic enzymes, and upregulation of iron import permeases (*feoA* and *feoB*), hemerythrin, *ideR*, flavodoxin and other flavoproteins. This assertion is supported by strong upregulation of flavodoxin in *C. beijerinckii*_mgsA+mgR (Fig. 5). Flavodoxins are electron-transfer proteins that participate in a wide variety cellular reactions (55). Interestingly, flavodoxins are prevalent in cells growing in iron-limited environments (55–57). In the present study, flavodoxin was the most upregulated gene. Furthermore, whereas 3 distinct Fe-S cluster assembly machinery have been characterized in bacteria, irrespective of the assembly type, cysteine desulfurase mediates the assembly of transient clusters on scaffold proteins and consequent relocation of pre-formed clusters to apo-proteins (58). It is plausible that down-regulation of Fe-S proteins and limited intracellular iron concentration within the cell elicited attendant downregulation of *csd* and ultimately, Fe-S cluster biosynthesis and assembly. This is possibly linked to downregulation of cysteine and methionine (sulfur bearing amino acids) uptake and metabolism, of which cysteine plays an important role in the anchoring of Fe-S clusters onto designated proteins (58).

**Fig 5.**
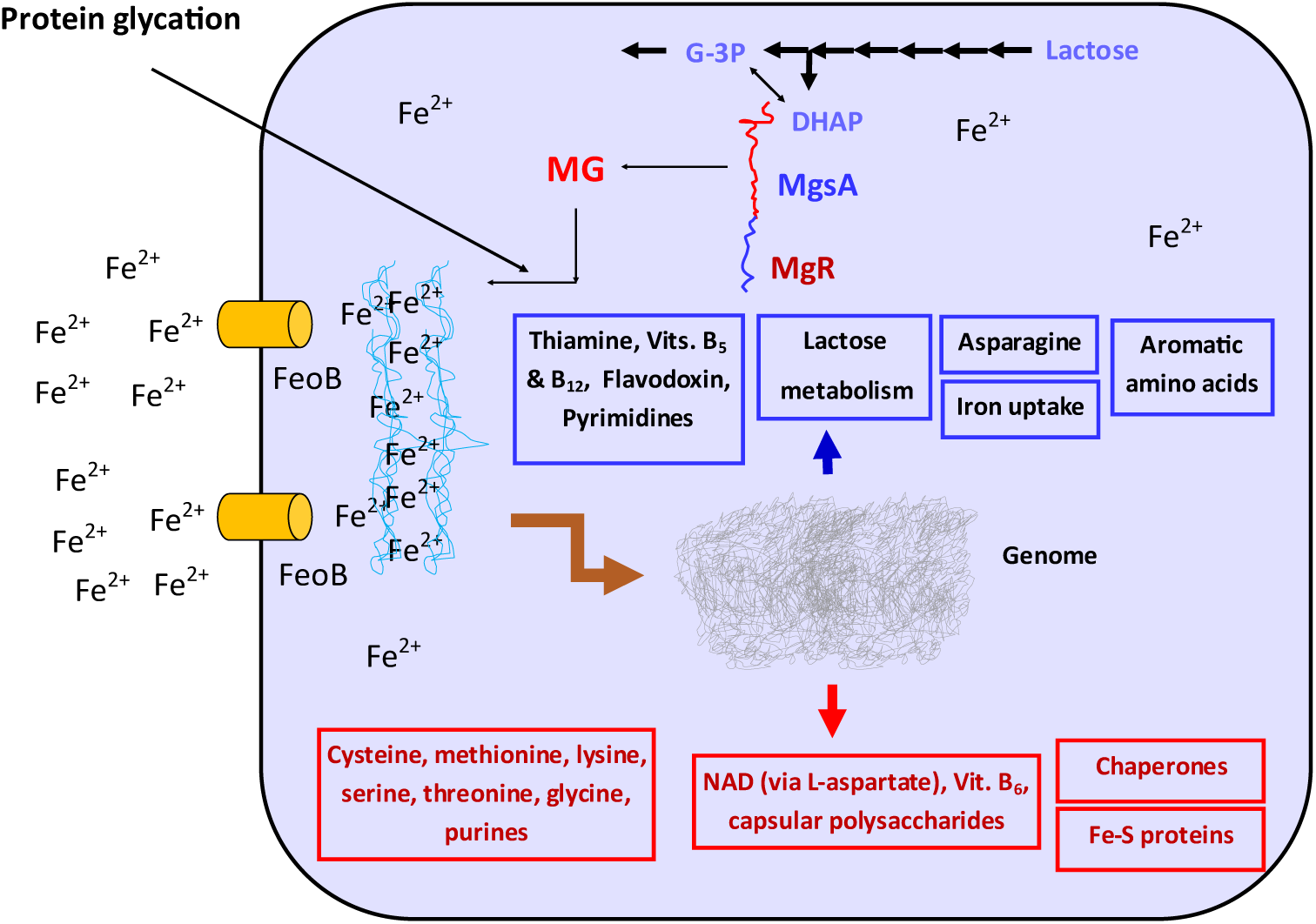
Summary of gene expression profile of lactose grown *C. beijerinckii*_mgsA+mgR relative to *C. beijerinckii*_p459, depicting select upregulated functions/pathways. G-3-P – Glyceraldehyde-3-phosphate; DHAP –Dihydroxyacetone phosphate. **↑** Upregulation; **↓** Downregulation.

The expression of genes involved in the uptake of purines, pyrimidines and peptides increased in *C. beijerinckii*_mgsA+mgR relative to *C. beijerinckii*_p459, whereas ammonium uptake was downregulated alongside biosynthesis of cysteine, methionine, lysine, asparagine, valine, glycine, serine, threonine, leucine, and isoleucine. Compared to ammonium, *C. beijerinckii* favors organic nitrogen sources, which might explain increased expression of a peptide uptake permease and concomitant downregulation of ammonium uptake. However, it is not clear what engendered this trend in *C. beijerinckii*_mgsA+mgR in comparison to *C. beijerinckii*_p459. Nonetheless, it is conceivable that greater reliance on organic nitrogen sources (peptides) by *C. beijerinckii*_mgsA+mgR, coupled with significantly improved lactose uptake and catabolism in this strain, led to improved growth and plausibly, enhanced solventogenesis when compared to *C. beijerinckii*_p459 (16). Furthermore, biosynthesis of purines and pyrimidines is ‘resource- and energy-intensive’, hence, salvaging these molecules from the immediate environment reduces cellular metabolic burden, which promotes growth.

Enhanced growth and cell viability can be manifested in the form of improved cell motility and downregulation of stress-responsive genes. Thus, the expression patterns of two-component signal transduction and cell motility genes in *C. beijerinckii*_mgsA+mgR clearly indicate overall greater cell viability and minimized stress on *C. beijerinckii*_mgsA+mgR, when compared to *C. beijerinckii*_p459. Conspicuously, the cell motility and signaling genes, *fliI* (flagellar protein export ATPase FliI), *flhF* (flagellar biosynthesis regulator FlhF), and *fliM* (flagellar motor switch protein FliM) were significantly upregulated in *C. beijerinckii*_mgsA+mgR. In fact, in total, 12 genes directly involved in flagellum biosynthesis, assembly and motor function were upregulated in *C. beijerinckii*_mgsA+mgR relative to *C. beijerinckii*_p459. Furthermore, among other genes, *manC* (Mannose-1-phosphate guanylyltransferase), *rfbA* (Glucose-1-phosphate thymidylyltransferase), *rfbB* (dTDP-glucose 4,6-dehydratase), *rfbC* (dTDP-4-dehydrorhamnose 3,5-epimerase), *murA* (UDP-N-acetylglucosamine 1-carboxyvinyltransferase), and *wbpA1* (UDP-glucose 6-dehydrogenase; Table S2) that take part in the biosynthesis of capsular polysaccharides exhibited reduced mRNA abundance in *C. beijerinckii*_mgsA+mgR. In addition, heat shock proteins and in some cases, sporulation genes, which typically signify exposure to a stressor were largely downregulated in *C. beijerinckii*_mgsA+mgR.

Capsular polysaccharides mediate biofilm formation and adherence of bacterial cells to surfaces (59). Thus, reduced mRNA abundance of capsular polysaccharide biosynthesis genes suggests that the cells of *C. beijerinckii*_mgsA+mgR are more discrete and motile. This notion is supported by upregulation of multiple signal transduction and cell motility genes in *C. beijerinckii*_mgsA+mgR (Table S2), coupled with the appearance of cells of *C. beijerinckii*_mgsA+mgR as largely discrete cells, whereas cells of *C. beijerinckii*_p459 showed greater clustering among cells. This can be ascribed to improved production of capsular polysaccharides in *C. beijerinckii*_p459 (Fig. 5). Remarkably, in addition to the flagella genes listed above, 38 genes involved in the signal transduction machinery (reception, processing and transmission of environmental cues) were upregulated in *C. beijerinckii*_mgsA+mgR, but not in *C. beijerinckii*_p459. Therefore, it can be deduced that *C. beijerinckii*_mgsA+mgR possesses a robust signal transduction machinery that likely favors nutrient acquisition and ultimately, growth and solventogenesis.

Despite its pivotal role under fermentative condition, *de novo* biosynthesis of NAD was downregulated in *C. beijerinckii*_mgsA+mgR. The protein product of *nadA* (quinolinate synthetase), a key step in the NAD *de novo* biosynthesis pathway via L-aspartate, is an Fe-S protein. Given considerable downregulation of Fe-S proteins in *C. beijerinckii*_mgsA+mgR, it is possible that the expression of *nadA* was depressed as a result of this broader trend (i.e., downregulation of Fe-S ptoteins). Further, *nadA* lies in an operon with *nadB* (L-aspartate oxidase) and *nadC* (nicotinate-nucleotide pyrophosphorylase/quinolinate phosphoribosyl transferase), both of which participate in *de novo* biosynthesis of NAD. Thus, downregulation of *nadA* (an Fe-S protein) likely contributed to downregulation of *nadB* and *nadC*. Despite the importance of NAD, *C. beijerinckii*_mgsA+mgR appears to exhibit a remodeled metabolic profile that circumvents the need for robust NAD biosynthesis. For instance, NADH dehydrogenase (Cbei_4112), which re-oxidizes NADH was upregulated 1.34-fold. At the time of sampling for RNA extraction (24 h), butanol concentrations—an outlet for NADH re-oxidation—were similar between *C. beijerinckii*_mgsA+mgR and *C. beijerinckii*_p459 (16). Concomitantly, Fe-only hydrogenase (Cbei_4110)—which is also involved in NADH re-oxidation was upregulated 1.25-fold. Since both NADH dehydrogenase and Fe-only hydrogenase are important players in NADH re-oxidation, the actions of both enzymes likely allow *C. beijerinckii*_mgsA+mgR to maximize available NAD via re-oxidation, thus mitigating reduced *de novo* biosynthesis. In fact, the growth and solvent profiles of *C. beijerinckii*_mgsA+mgR relative to *C. beijerinckii*_p459 suggest that the prevailing metabolic machinery of the former is more efficient than that of the latter. Thus, addition of 2 g/L L-aspartate may have upregulated the L-aspartate end of NAD *de novo* biosynthesis via L-aspartate, possibly at the expense of another salient nexus in the metabolic network, which in turn, disrupted the metabolic sequence in this strain, thereby leading to severely diminished growth, ABE production, and acid re-assimilation (Figs. 2 & 3). The results suggest that *de novo* biosynthesis of NAD in *C. beijerinckii*_mgsA+mgR undergoing possible MG-induced intracellular iron limitation likely steals “scarce” iron and perhaps, other metabolic resources away from growth and solvent production.

Whereas the genes involved the metabolism of cysteine, methionine, lysine, asparagine, glycine, serine, threonine, valine leucine, and isoleucine were downregulated, genes associated with the synthesis of aromatic amino acids (particularly, tryptophan) and arginine were upregulated. As mentioned earlier, downregulation of cysteine biosynthesis is ascribable to down-regulation of Fe-S proteins. On the other hand, reduced expression of genes associated with the metabolism of asparagine, methionine, threonine, and lysine may be connected to the aforementioned shift of metabolic processes away from L-aspartate. Indeed, asparagine, methionine, threonine, and lysine are all aspartate-derived amino acids (61). It appears therefore, that biochemical reactions downstream of aspartate—and possibly upstream—were largely depressed in *C. beijerinckii*_mgsA+mgR. This effect is clearly not as a result of aspartate limitation because, growth in an L-aspartate-replete medium led to severely impaired growth and butanol production. Downregulation of *asd* (aspartate-semialdehyde dehydrogenase), which catalyzes a key step in the biosynthesis of aspartate and ultimately, biosynthesis of methionine, lysine, and threonine from aspartate (62), is a strong indication that the production of aspartate and its derivatives were actively reduced in *C. beijerinckii*_mgsA+mgR during lactose fermentation. Remarkably, aspartate is involved in asparagine, isoleucine, methionine, threonine, pyrimidines, NAD, and vitamin B_5_ biosynthesis and as a nitrogen donor for arginine and purine syntheses (63). Among these functions, biosyntheses of arginine, vitamin B_5_, and pyrimidines were upregulated in lactose-grown *C. beijerinckii*_mgsA+mgR, albeit inexplicable. However, it is important to highlight that aspartate is also a precursor of purine and pyrimidine, as is glycine— whose metabolism was equally downregulated—in pyrimidine synthesis. Thus, reduced expression of aspartate and glycine metabolic genes, may free up more of both precursors for pyrimidine biosynthesis, which was upregulated in *C. beijerinckii*_mgsA+mgR. Therefore, enhanced purine uptake and biosynthesis of pyrimidines likely combined to promote nucleotide biosynthesis and ultimately, replication in *C. beijerinckii*_mgsA+mgR, when compared to *C. beijerinckii*_p459. This may explain the 35% higher growth observed with *C. beijerinckii*_mgsA+mgR relative *C. beijerinckii*_p459 (16).

The factors above do not explain increased expression of genes involved in the production of aromatic amino acids. The protein product of *aroH* (phospho-2-dehydro-3-deoxyheptonate aldolase)—which was upregulated in *C. beijerinckii*_mgsA+mgR—catalyzes the first of 7 steps leading to biosynthesis of chorismate, a precursor of all three aromatic amino acids, *p*–amino butyric acid, folate cofactors, ubiquinone and menaquinone (61, 64). Concomitantly, only *trpC* (indole-3-glycerol-phosphate synthase) and *trpE* (anthranilate synthase component I), both of which are involved in tryptophan biosynthesis were further upregulated in this pathway, which might indicate that tryptophan was the major biosynthesis target in the aromatic amino acid pathway. Asides aspartate, tryptophan is a precursor of quinolinate in an alternative arm of the *de novo* NAD biosynthesis pathway. Thus, we speculate that increased expression of tryptophan biosynthesis genes might represent an alternative route for NAD production in *C. beijerinckii*_mgsA+mgR. Since *asd* was downregulated in *C. beijerinckii*_mgsA+mgR, the eventual diversion of L-aspartate semialdehyde would broadly limit L-aspartate-dependent biochemical reactions, most of which were downregulated in this study. We are currently using RNA sequencing and isotope labeling to characterize MG-challenged cultures of *C. beijerinckii*, in an effort to better delineate the response of this organism to MG. We expect that the result will provide additional insight that will aid metabolic engineering of a robust strain capable of efficient biosynthesis 1,2-PD via the MG bypass. Also, the result will likely enhance further engineering of *C. beijerinckii*_mgsA+mgR for improved butanol production.

## CONCLUSION

In this study, we sought to understand the basis for enhanced production of butanol by *C. beijerinckii*_mgsA+mgR, a recombinant strain of *C. beijerinckii* heterologously expressing *mgsA* and *mgR* from *C. pasteurianum*. The mRNA profiles suggest reduced expression of L-aspartate-dependent enzymes and their downstream genes including *de novo* NAD biosynthesis. Conversely, the expression profiles suggest that *C. beijerinckii*_mgsA+mgR might deploy the tryptophan route to supplement NAD production. In parallel, genes involved in iron import were significantly upregulated alongside downregulation of a large number of gene encoding Fe-S proteins. This is indicative of intracellular ferrous iron limitation, possibly due to iron complexing with methylglyoxal glycated proteins. Upregulation of flavodoxin and other flavoproteins further supports the notion that ferrous iron is limited in *C. beijerinckii*_mgsA+mgR. Increased expression of genes involved in purine and pyrimidine import and pyrimidine biosynthesis and lactose uptake and utilization are in line with the enhanced growth and solvent production previously observed with *C. beijerinckii*_mgsA+mgR.

## Acknowledgements

This research was financially supported by awards from USDA-National Institute of Food and Agriculture Hatch Program (grant no. WIS04018), the Dairy Innovation Hub (grant no. AAK5475), and Dairy Business Innovation Alliance (DBIA) to V.C.U.

